# The astrovirus N-terminal nonstructural protein anchors replication complexes to the perinuclear ER membranes

**DOI:** 10.1101/2024.01.09.574783

**Authors:** Hashim Ali, David Noyvert, Jacqueline Hankinson, Gemma Lindsey, Valeria Lulla

## Abstract

An essential aspect of viral replication is the anchoring of the replication complex (RC) to cellular membranes. Positive-sense RNA viruses employ diverse strategies, including co-translational membrane targeting through signal peptides and co-opting cellular membrane trafficking components. Often, N-terminal nonstructural proteins play a crucial role in linking the RC to membranes, facilitating the early association of the replication machinery. Astroviruses utilize a polyprotein strategy to synthesize nonstructural proteins, relying on subsequent processing to form replication-competent complexes. In this study, we provide evidence for the perinuclear ER membrane association of RCs in five distinct human astrovirus strains. Using tagged recombinant classical human astrovirus 1 and neurotropic MLB2 strains, we establish that the N-terminal domain guides the ER membrane association. Through mutational analysis of the N-terminal domain in replicon and reverse genetics systems, we identified di-arginine motifs responsible for the perinuclear ER retention and formation of functional RCs. Our findings highlight the intricate virus-ER interaction mechanism employed by astroviruses, potentially leading to the development of novel antiviral intervention strategies.

**Author Summary:** Human astroviruses are a significant cause of acute gastroenteritis, accounting for up to 9% of cases in young children. Immunocompromised individuals and infants experience more critical symptoms, such as severe and persistent diarrhea, as well as sporadic systemic and even fatal diseases. To date, no drugs have been developed to protect against astrovirus infection. Our study provides the first evidence that the integrity of the N-terminal domain of nsP1a is essential for establishing early replication. Central to this process, the di-arginine motifs in the N-terminal domain are responsible for ER retention, the formation of functional replication complexes, and viral replication. Therefore, selectively targeting N-terminal domain-mediated ER retention could be a promising therapeutic strategy to effectively control astrovirus infection.

## Introduction

Astroviruses are a family of small non-enveloped (+)ssRNA viruses that infect a wide range of mammalian and avian species. Humans are susceptible to the classical (HAstV1-8) and neurotropic (VA/HMO and MLB) clades of astroviruses, which cause mild to severe diseases depending on the age and health of an individual ^1,2^. Despite the high zoonotic potential and non-gastrointestinal tropisms ^2–4^, there are still significant gaps in our understanding of the biology of human astroviruses and disease progression. Besides, astrovirus infections are often under-reported despite their high prevalence ^5^. Currently, no vaccines or drugs against astroviruses are available. Therefore, it is important to investigate their replication mechanisms to effectively control future outbreaks.

The astrovirus genome is approximately 6.8–7.9 kilobases (kb) in length, linked to genome-linked viral protein (VPg) at the 5′ end and polyadenylated at the 3′ untranslated region ^1,6^, with four open reading frames (ORFs): ORF1a, ORF1b, ORFX and ORF2 ^7,8^. Both ORF2 and ORFX are translated from subgenomic RNA and encode structural proteins and viroporin, respectively ^7,9^. ORF1b encodes the RNA-dependent RNA polymerase (RdRp), translated following ribosomal frameshifting in the overlapping region between ORF1a and ORF1ab ^1,10^. ORF1a encodes a large nonstructural polyprotein (nsP1a), which is processed into several proteins, including the N-terminal protein encoding putative RNA helicase, serine protease, VPg, and p20 protein ^1^. The exact number of nonstructural proteins and polyprotein processing rules are yet to be determined.

Replication of (+)ssRNA viruses requires RdRp and other nonstructural proteins that form the viral replication complex. Viruses exploit host membranes to assemble replication complexes (RCs), protect dsRNA intermediates, and segregate replicating RNAs from translating RNAs ^11–13^. In astroviruses, the components of the replication complex contain RdRp, protease, and VPg – three well-characterized enzymatic units ^6,10,14^. However, membrane-anchoring and retention strategies are still unknown. The predicted transmembrane (TM) domain located at the N-terminal part of the astrovirus nonstructural protein suggests its involvement in membrane association. Recently, ER-derived membranes were implicated in the formation of double-membrane vesicles during astrovirus infection ^15^. However, the exact location, ER membrane specificity, and viral proteins responsible for this remain to be characterized. In addition to the TM domain, the N-terminal domain of the astrovirus polyprotein contains a putative RNA helicase motif ^1^. However, its poor conservation and incomplete motif integrity raise questions regarding the functional significance of the proposed helicase.

Here, we demonstrate the function of the N-terminal protein in two astrovirus genotypes, HAstV1 and MLB2. This small membrane protein drives the formation of replication complexes in tight association with perinuclear ER membranes, which is a key feature of RNA replication in several astroviruses (HAstV1, HAstV4, MLB1, MLB2, and VA1). We also found that ER retention and function of the astrovirus replication complex are dependent on the di-arginine motif located in the N-terminal protein. In the future, an improved understanding of astrovirus replication complex formation may represent a promising therapeutic strategy for controlling astrovirus-associated infections in young children, immunocompromised individuals, and farm animals.

## Results

### The putative helicase domain is not conserved and is dispensable for HAstV1 replication

RNA viruses harbor several cis-acting elements that play essential roles in viral RNA replication, translation and assembly of virions ^16^. During viral genome replication, these structured RNA elements require RNA helicases or chaperones to facilitate the unwinding and remodeling of double-stranded RNA structures. Helicases use energy derived from NTP hydrolysis to catalyze unwinding and contain several conserved motifs, including NTP-binding Walker A and Walker B motifs ^17^. Several RNA viruses encode NTPase/RNA helicases, which assist in the unwinding of dsRNA replicative intermediates during virus replication ^18^. Interestingly, astrovirus genomes do not have features that can be attributed to a fully functional NTPase/helicase. The only Walker A-like motif can be found in classic human astroviruses, but not in neurotropic MLB and VA genotypes (Fig 1A), suggesting poor conservation and questioning the presence of virus-encoded helicases in astroviruses. In addition, this domain is predicted to be attached to the membrane and separated from the rest of the replication module (Fig 1B), which is an unusual localization for RNA-processing enzymes. To address this question, the putative Walker A motif in HAstV1 was mutated from GKT to GAT in the context of replicons and infectious viruses. In the replicon system, a minor reduction in activity was observed (Fig 1C), and in the GKT-to-GAT recombinant virus, no significant differences were observed (Fig 1D), whereas mutation of functional GKT/GKS motifs in other viruses is usually lethal ^19,20^. This further indicates that the GKT motif is unlikely to play a functionally important role in the HAstV1 replication. In addition, several viral NTPases/helicases, including enterovirus 2C^ATPase^ ^21^, are inhibited by guanidine hydrochloride (GuHCl), resulting in NTPase-specific inhibition of replication ^22^. Using HAstV1-based and enterovirus-based replicon systems, we examined the ability of GuHCl to inhibit replication. Unlike previously characterized inhibition in enteroviruses, astrovirus replication was not strongly affected by GuHCl treatment (Fig 1E-F). The 20-30% decrease in astrovirus replicon activity is at least partially caused by the decrease in cell confluency in the presence of GuHCl at later time points (Fig 1E), whereas enterovirus replication was efficiently inhibited to ∼0.1% (Fig 1F), further confirming that GuHCl-sensitive replicase components are absent in astroviruses. Considering the poor conservation (Fig 1A) and lack of functional effect (Fig 1C-E) of the putative Walker A motif in HAstV1-based assays, it is plausible to suggest that cellular helicases can be recruited by astrovirus replication complexes (RCs) as an alternative strategy utilized by several virus families ^23^. Therefore, the remaining GKT motif may represent an evolutionarily lost NTPase/helicase rather than a functional unit within the astrovirus genome.

**Fig 1.**
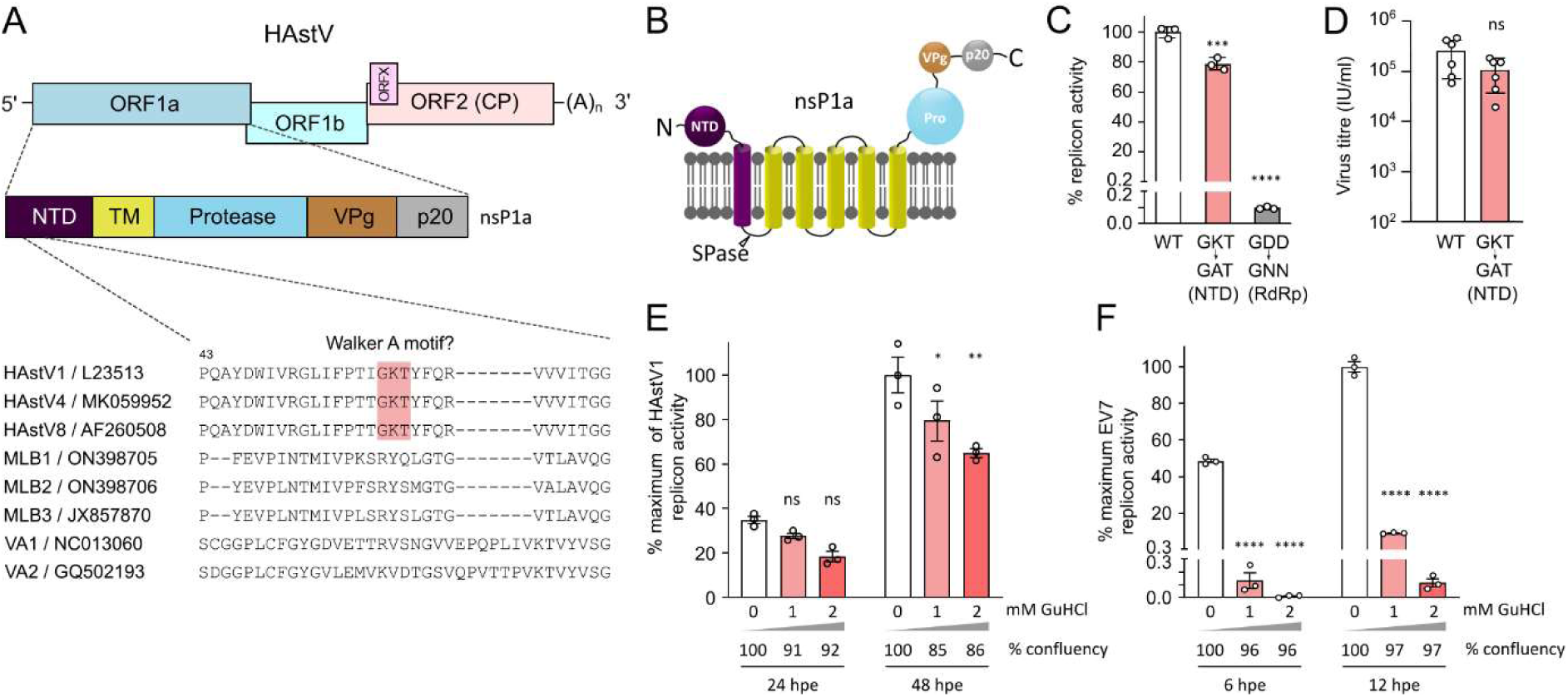
Analysis of the role of the N-terminal putative helicase in astrovirus replication. **(A)** Schematic representation of the astrovirus genome, location of N-terminal domain (NTD), and putative Walker A motif, aligned for several astrovirus genomes. **(B)** Predicted domain organization of the astrovirus nonstructural polyprotein. SPase, signal peptidase; Pro, protease domain; VPg, viral protein genome-linked. C- and N-terminal ends of polyprotein are indicated. **(C)** The activity of HAstV1-based Renilla luciferase-expressing replicon was measured in Huh7.5.1 cells at 24 hpt. **(D)** The virus titer was measured in Caco2 cells after electroporation of BSR cells with *in vitro* transcribed T7 HAstV1 RNA. **(E-F)** The effect of guanidine hydrochloride (GuHCl) on the activity of mCherry-expressing astrovirus (E) and enterovirus (F) replicons. The mean cell confluence is provided for each time point. Data are mean ± SEM. **** *p* < 0.0001, *** *p* < 0.001, ** *p* < 0.01, * *p* < 0.1, ns, nonsignificant, using one-way ANOVA (C), Student’s t-test (D) or two-way ANOVA (E-F), against WT (C-D) or untreated control (E-F).

### Role, accumulation and localization of NTD in astrovirus replication

To investigate the role(s) of the NTD in virus replication, we engineered HA-tagged astroviruses using classical HAstV1 ^7^ and neurotropic MLB2 ^24^ reverse genetics systems by placing an HA-tag sequence in the predicted disordered region between the folded N-terminal part of the protein and the first transmembrane helix. If N-terminal cleavage occurs at the predicted signal peptidase (SPase) cleavage site, the molecular weight of the N-terminal HA-tagged astrovirus protein is predicted to be 21-22 kDa (Fig 2A). MLB2-HA virus was successfully rescued, and growth kinetics was assessed in Huh7.5.1 cells, displaying a minor delay in growth (Fig 2B), which is common for tagged viruses with small genomes. The rescue of HAstV1-HA was successful on transfection but not on passaging (Fig 2C). This could indicate a defect at later stages of infection in Caco2 cells. The accumulation of 14 kDa HA-tagged product was detected in HA-tagged but not in wt astrovirus-infected cells (Fig 2D-E), suggesting potential N-terminal cleavage of NTD or altered mobility. Capsid accumulation showed a slight delay in HA-tagged MLB2 samples when compared to the wt virus, consistent with the delay in virus growth (Fig 2D). Next, we analyzed the same viral products using immunofluorescence of the virus-infected cells at 24 hpi. Interestingly, we observed a distinct perinuclear localization of HA-tagged NTD for both viruses, whereas the capsid protein was dispersed throughout the cytoplasm (Fig 2F-G), suggesting different sites of replication and packaging in astrovirus-infected cells. Notably, the cytoplasmic distribution of the capsid was more evident in Huh7.5.1 cells due to their larger cytoplasmic space (Fig 2F).

**Fig 2.**
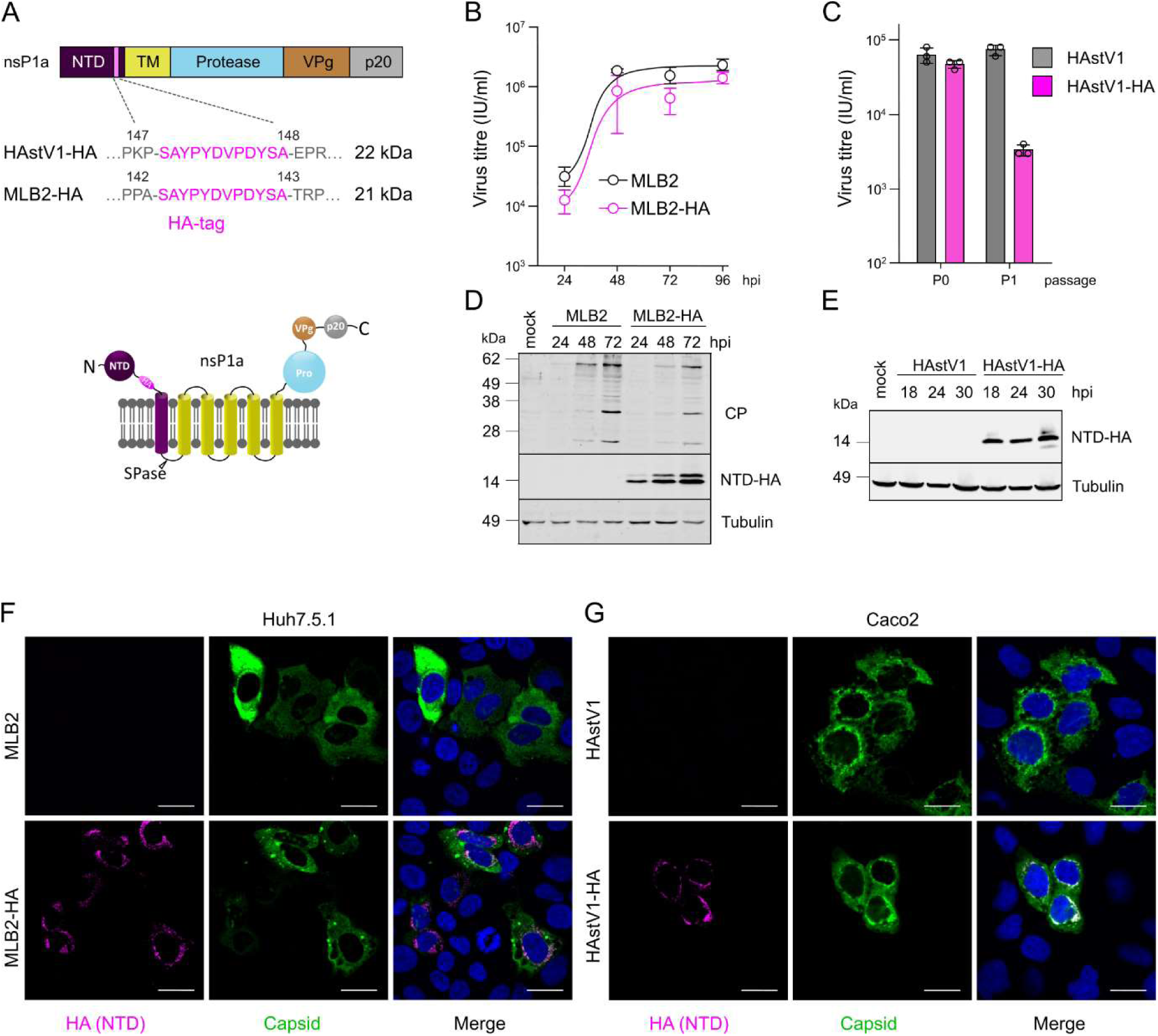
Design and characterization of HA-tagged astroviruses. **(A)** Schematic representation of astrovirus ORF1a polyprotein with HA-tagged NTD domain. The amino acid positions are indicated for each astrovirus strain (top panel). Predicted membrane topology for HA-tag within nsP1a polyprotein (bottom panel). Predicted molecular weight is provided for HAstV1-HA and MLB2-HA SPase-cleaved N-terminal protein. **(B)** Multistep growth curves of MLB2 and MLB2-HA viruses in Huh7.5.1 cells. Cells were infected at an MOI 0.1 (passage 2), and virus titer was measured from the extracellular fractions in triplicates. Data are mean ± SEM. **(C)** Titers of rescued (passage 0) and passaged (passage 1 in Caco2 cells) HAstV1 and HAstV1-HA viruses. **(D)** Huh7.5.1 cells were infected at an MOI 0.1 (passage 2), harvested at indicated hpi and analyzed by western blotting with anti-CP and anti-HA antibodies. **(E)** Caco2 cells were infected at an MOI 1 (passage 0), harvested at indicated hpi and analyzed by western blotting with anti-HA antibodies. **(F-G)** Representative confocal images of fixed and permeabilized cells visualized for MLB2 (F) or HAstV1 (G) CP (green) and HA-tag (magenta). Nuclei were stained with Hoechst (blue). Scale bars are 25 µm.

### Astroviruses replicate in the endoplasmic reticulum, close to the nuclear periphery

Similar to other (+)ssRNA viruses, human astroviruses replicate in the host endoplasmic reticulum (ER) by forming double-membrane vesicles ^15^. However, the mode of RC formation and its retention within the ER membranes still needs to be determined. To systematically investigate the cellular localization of astrovirus replication sites, we used five available astrovirus strains (HAstV1, HAstV4, MLB1, MLB2 and VA1) and infected two different cell lines (Huh7.5.1 and Caco2) that support selected astrovirus replication and spread. Staining with anti-dsRNA antibody, a hallmark of (+)ssRNA replication sites, revealed perinuclear localization of RCs in Huh7.5.1 (Fig 3A) and Caco2 (Fig 3B) cells. The dsRNA-specific signal overlapped with ER-specific staining only very close to the nuclear periphery, but not further in the cytoplasm in all tested astrovirus-infected cells (Fig 3). Co-staining with lamin, a component of the nuclear membrane, revealed partial co-localization (Fig 4), confirming the preferential perinuclear localization of the astrovirus RCs. These results suggest the specific targeting of RCs to the perinuclear ER membranes and the possible involvement of NTD in this process (Fig 2F-G).

**Fig 3.**
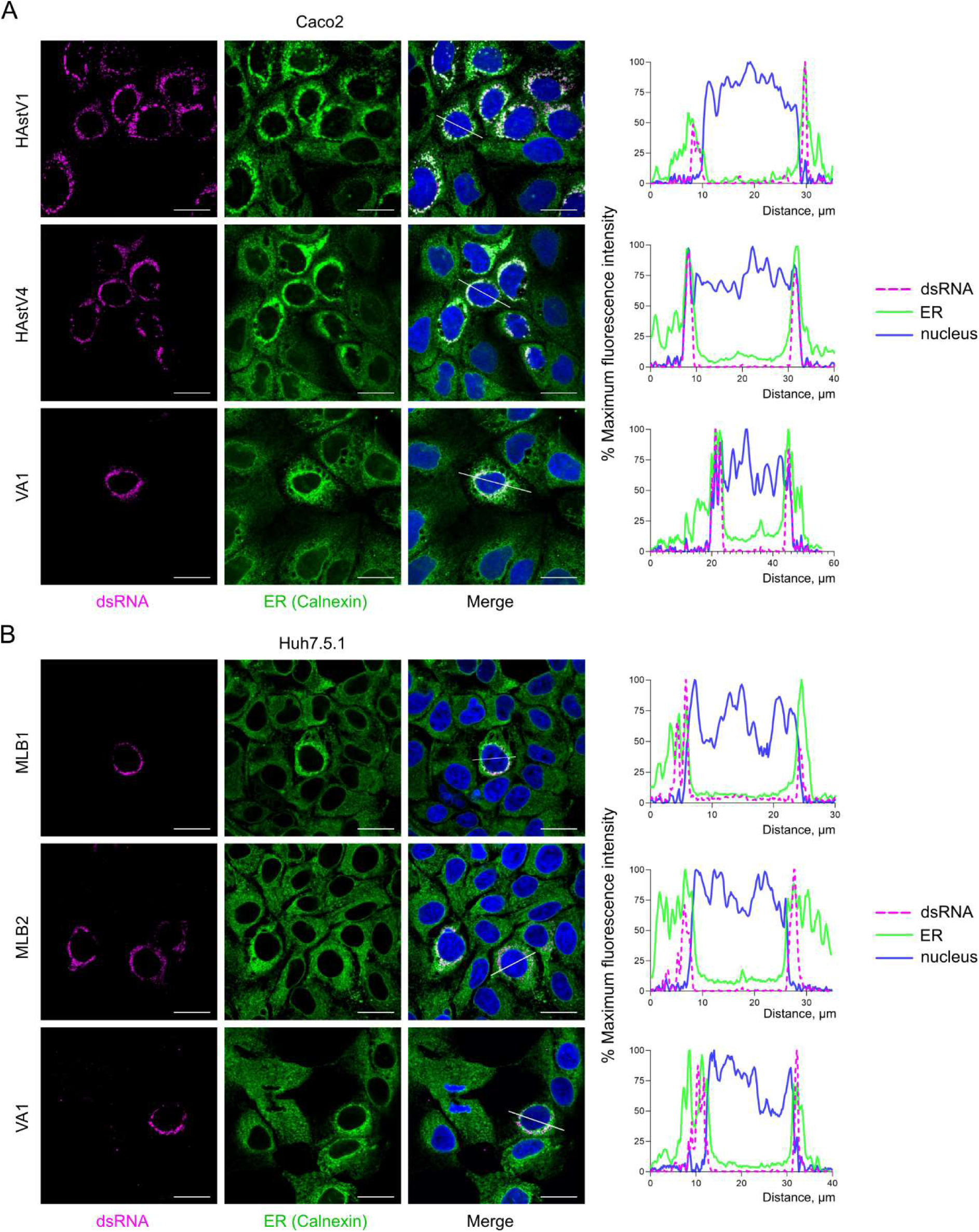
Astrovirus replication sites are localized in the perinuclear ER membranes. **(A)** Caco2 cells were infected with HAstV1, HAstV4 and VA1 astroviruses. (**B**) Huh7.5.1 cells were infected with MLB1, MLB2 and VA1 astroviruses. (A-B) Representative confocal images of fixed and permeabilized cells visualized for dsRNA (magenta) and ER (calnexin, green). Nuclei were stained with Hoechst (blue). Scale bars are 25 µm. Intensity profiles of dsRNA (magenta), calnexin (green) and nuclear staining (blue) were obtained using ImageJ software, along a straight line shown on the merged image crossing the representative cell.

**Fig 4.**
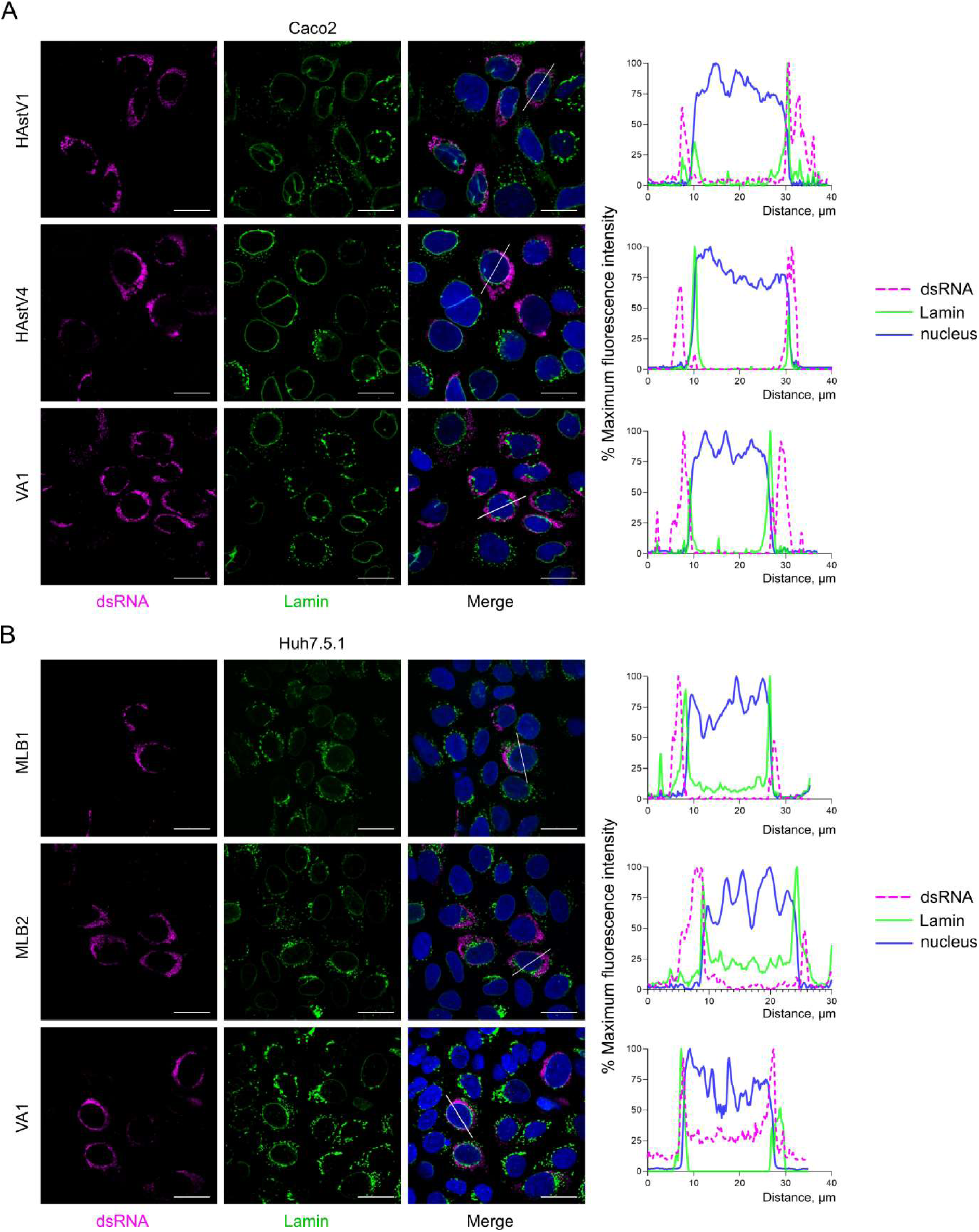
Astrovirus replication sites demonstrate perinuclear localization. **(A)** Caco2 cells were infected with HAstV1, HAstV4 and VA1 astroviruses. (**B**) Huh7.5.1 cells were infected with MLB1, MLB2 and VA1 astroviruses. (A-B) Representative confocal images of fixed and permeabilized cells visualized for dsRNA (magenta) and nuclear lamin (green). Nuclei were stained with Hoechst (blue). Scale bars are 25 µm. Intensity profiles of dsRNA (magenta), lamin (green) and nuclear staining (blue) were obtained using ImageJ software, along a straight line shown on the merged image crossing the representative cell.

### The NTD is co-localized with replicating RNA during astrovirus infection

To investigate the localization of NTD during astrovirus infection, Caco2 and Huh7.5.1 cells were infected with HAstV1-HA and MLB2-HA, respectively. As expected, a nearly complete overlap between dsRNA and NTD was observed (Fig 5), confirming the previous observations. Consistent with the staining obtained with wt astrovirus strains (Fig 3-4), HA-tagged NTD-specific localization followed the same overlapping perinuclear ER- and lamin-specific pattern (Fig 5). Overall, these results confirmed the association of the astrovirus NTD with RNA RCs, raising an important question: what drives the NTD to the perinuclear ER membranes and how does this affect astrovirus replication?

**Fig 5.**
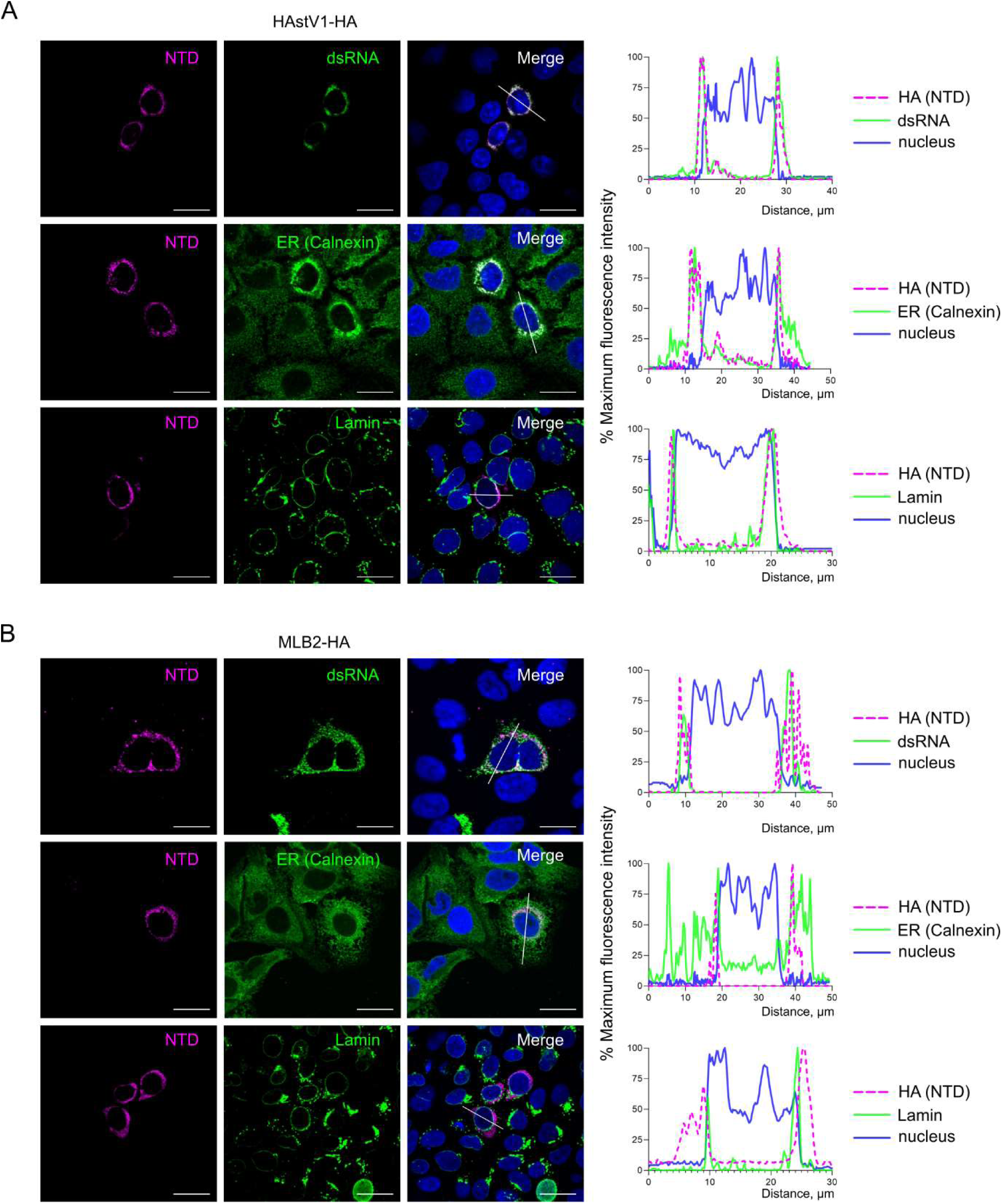
Astrovirus NTD is linked to RNA replication sites. **(A)** Caco2 cells were infected with HAstV1-HA. (**B**) Huh7.5.1 cells were infected with MLB2-HA. (A-B) Representative confocal images of fixed and permeabilized cells visualized for HA-tag (NTD, magenta), dsRNA (green), ER (calnexin, green), and lamin (green). Nuclei were stained with Hoechst (blue). Scale bars are 25 µm. Intensity profiles of NTD (magenta), dsRNA (green), ER (green), lamin (green), and nuclear staining (blue) were obtained using ImageJ software, along a straight line shown on the merged image crossing the representative cell.

### Di-arginine motifs in NTD are responsible for perinuclear ER localization and astrovirus replication

ER targeting of RCs in (+)ssRNA viruses can be achieved through various mechanisms ^25^. In astroviruses, the NTD, followed by transmembrane helices, represents a unique combination to target the nonstructural polyprotein to the correct position within the infected cell. We explored the power of molecular mimicry used by many viruses ^26^ to identify the potential residues responsible for ER targeting that were predicted using the ELM web server (http://elm.eu.org/) ^27^. The search revealed the presence of conserved di-arginine motifs (Fig 6A), which were defined by two consecutive arginine residues (RR) or with a single residue insertion (RXR). This motif is characteristic of several membrane proteins with ER localization and allows for correct folding and membrane association. The functional motif needs to be exposed to the cytoplasm and requires distinct proximity to the TM region, thus fulfilling all criteria for a predicted astrovirus nonstructural polyprotein (Fig 1B, 6A) ^28^.

**Fig 6.**
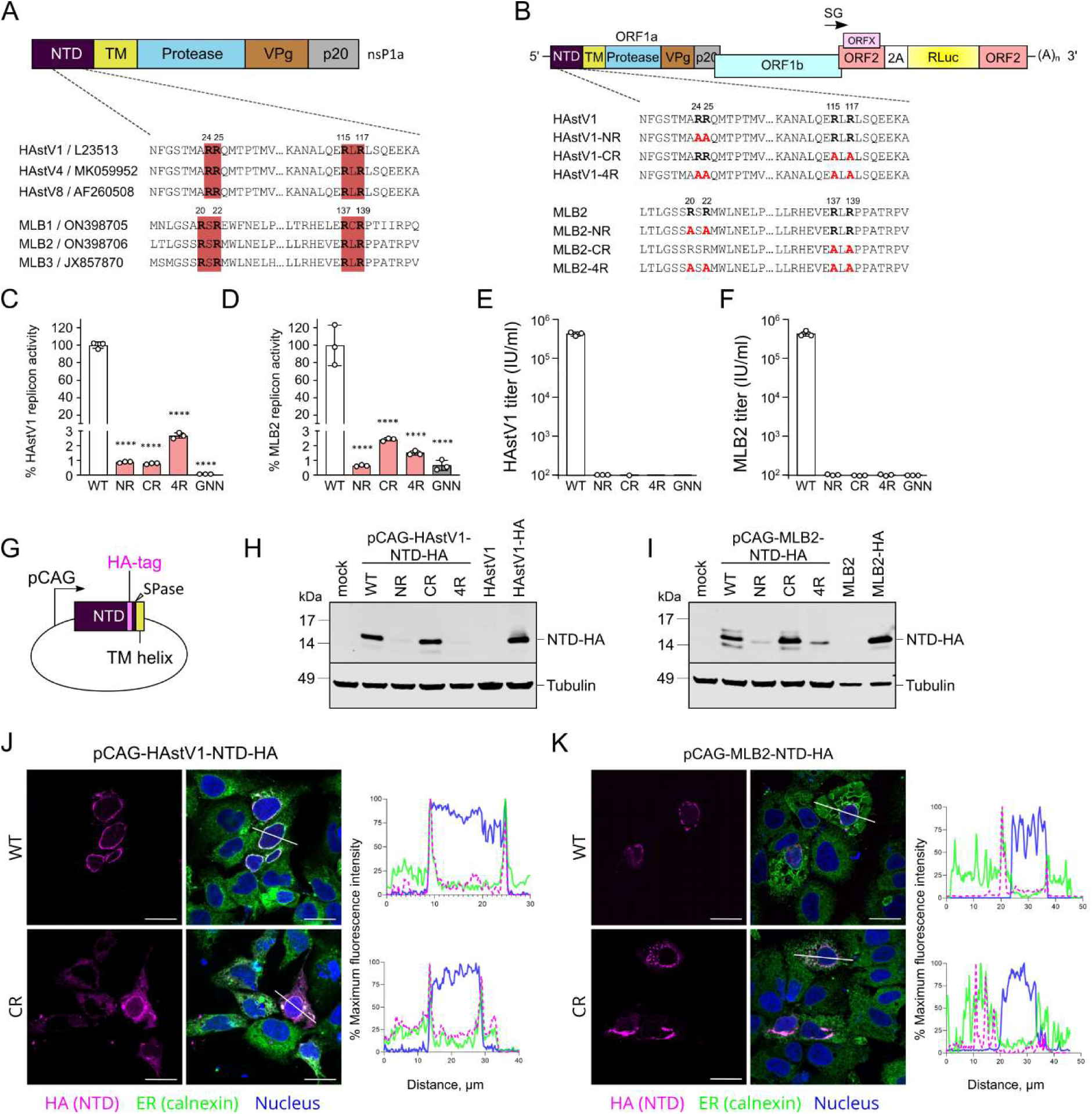
NTD-specific di-arginine motifs anchor astrovirus RC to perinuclear ER membranes and are required for virus replication. **(A)** Schematic representation of astrovirus nsP1a polyprotein with di-arginine motifs. The amino acid positions are indicated for each astrovirus strain. (**B**) Schematic representation of astrovirus replicon with introduced R-to-A mutations. (**C-D**) Relative luciferase activities of HAstV1 (C) and MLB2 (D) based replicons in Huh7.5.1 cells at 24 hours post-transfection. Data are mean ± SD. **** *p* < 0.0001, using one-way ANOVA test against WT. (**E-F**) Titers (infectious units per ml) of recombinant viruses after RNA electroporation of Huh7.5.1 cells followed by titration on Caco2 (HAstV1, E) or Huh7.5.1 (MLB2, F) cells. (**G**) Schematic representation of CAG promoter containing mammalian expression vector used to over-express HA-tagged NTD in Huh7.5.1 cells. (**H-I**) Huh7.5.1 cells were electroporated with pCAG plasmids expressing wt and mutant NTD-HA and analyzed by western blotting with anti-HA and anti-tubulin antibodies. Lysates obtained from infected cells were used as a control. (**J-K**) Huh7.5.1 cells were electroporated with pCAG plasmids expressing wt and mutant NTD-HA and analyzed by immunofluorescence. Representative confocal images of fixed and permeabilized cells visualized for HA-tag (NTD, magenta) and ER (calnexin, green). Nuclei were stained with Hoechst (blue). Scale bars are 25 µm. Intensity profiles of NTD (magenta), ER (green) and nuclear staining (blue) were obtained using ImageJ software, along a straight line shown on the merged image crossing the representative cell.

To investigate the function of the di-arginine motifs of NTD in the virus life cycle, arginine-to-alanine mutations were introduced into replicons and full-length infectious clones of HAstV1 and MLB2 astroviruses (Fig 6B). Replication was drastically decreased in all mutant replicons (Fig 6C-D) and all di-arginine mutant viruses were not viable (Fig 6E-F), confirming the critical role of both di-arginine motifs in the virus life cycle.

To link the replication defect to the specific ER localization performed by the NTD, the same subset of mutations (Fig 6B) was introduced into a mammalian expression vector that encodes only 187 (MLB2) or 190 (HAstV1) amino acid residues of the HA-tagged NTD (Fig 6G). All NTD variants containing mutated N-terminal di-arginine motifs showed reduced amounts of protein (Fig 6H-I), suggesting that the stability of the protein is dictated by the correct ER targeting mediated by the N-terminal di-arginine motif.

The perinuclear localization of overexpressed wt NTD-HA (Fig 6J-K, top panels) recapitulated the NTD-HA localization in virus-infected cells (Fig 5), suggesting the role of this domain in the correct anchoring of astrovirus RCs. Perinuclear ER staining was drastically altered in CR arginine-to-alanine mutants, resulting in a diffuse cytoplasmic pattern in cells expressing both HAstV1 and MLB2 NTD-HA protein, despite the TM helix and N-terminal di-arginine motif remaining unmodified (Fig 6J-K).

Taken together, these results provide evidence for the distinct features of N- and C-terminal di-arginine motifs and the functional role of the NTD in the membrane anchoring of astrovirus RC, which is a prerequisite for efficient replication.

## Discussion

In this study, we investigated the role of the astrovirus N-terminal protein in ER membrane tethering and the formation of functional RCs. We demonstrate that the putative helicase is unlikely to be a functional unit within astrovirus RCs. Instead, the di-arginine signature-driven ER membrane localization and replication represents a key role of the N-terminal protein in the astrovirus replication cycle.

Numerous positive-sense RNA viruses encode RNA helicases, while others depend on cellular counterparts in their absence ^18,23^. Interestingly, astrovirus genomes encode a single Walker A-like motif that can be found in classic human astroviruses but not in the newly emerging human (Fig 6A) and avian astroviruses ^29^. Consistent with poor conservation, the functional role of the GKT motif was not detrimental to HAstV1 replication and life cycle (Fig 1). The absence of evidence of a functional helicase in the genome of astroviruses suggests dependence on host proteins with NTPase/helicase activity. Interestingly, proteomic analysis of HAstV8-infected Caco2 cells showed enrichment of membrane-only fractions with the cellular RNA helicase DDX23, detected alongside viral RdRp and protease – ultimate components of RCs ^30^. Additionally, siRNA-mediated depletion of DDX23 significantly decreased virus replication ^30^, indicating the host helicase dependence of astrovirus replication. In a recent transcriptomic analysis of HAstV1-infected Caco2 cells, cellular helicases HELZ2 and DDX58 transcripts were found to be upregulated in infected cells ^15^, providing more candidates for the host helicases that can be involved in the RNA replication process.

The correct formation of (+)ssRNA virus RCs is mediated by the recruitment of cellular membranes that prevent immune detection of viral RNA, separate the processes of replication and translation, and increase the local concentration of active replication components, both viral and cellular ^11–13^. Viruses employ a range of strategies to ensure the correct association of replication-competent virus-host replication machinery. Co-opting the ER for this purpose has been described for numerous RNA viruses including the *Flaviridae*, *Coronaviridae*, *Picornaviridae* ^31^ families, and has also suggested for astroviruses ^15,30,32^. We confirm this localization and identify perinuclear ER membranes as a preferential RNA replication site (Fig 3), similar to several other RNA viruses ^33–35^.

The involvement of the N-terminal part of the nonstructural polyprotein in the positioning of the whole replication complex is a convenient strategy to ensure translocation across the ER membrane which can be mediated by the N-terminal signal/sorting peptide ^36^. In astroviruses, the N-terminal protein, followed by transmembrane helices, is ideally positioned to target the nonstructural polyprotein to cellular membranes. Of the multiple strategies of ER membrane integration ^36,37^, we found that astroviruses use di-arginine-based ER-sorting motifs ^38^ to anchor and assemble RCs. This strategy is used by several mammalian (lip35, GABA_B_, Kir6.2)^37^, plant (AtGCSI)^39^, and viral (hepatitis B virus S)^40^ proteins for membrane trafficking ^37^. We demonstrated that two predicted pairs of N-terminal di-arginine motifs are essential for virus replication and responsible for perinuclear ER localization of the N-terminal protein (Fig 6,7). This links two crucial functions together: the correct localization of the RC components and their activity. As many viruses exploit the ER during infection, pharmacological strategies aiming at disrupting virus-ER traffic/interactions or ER morphogenesis should, in principle, lead to the generation of broad-spectrum antiviral targets.

**Fig 7.**
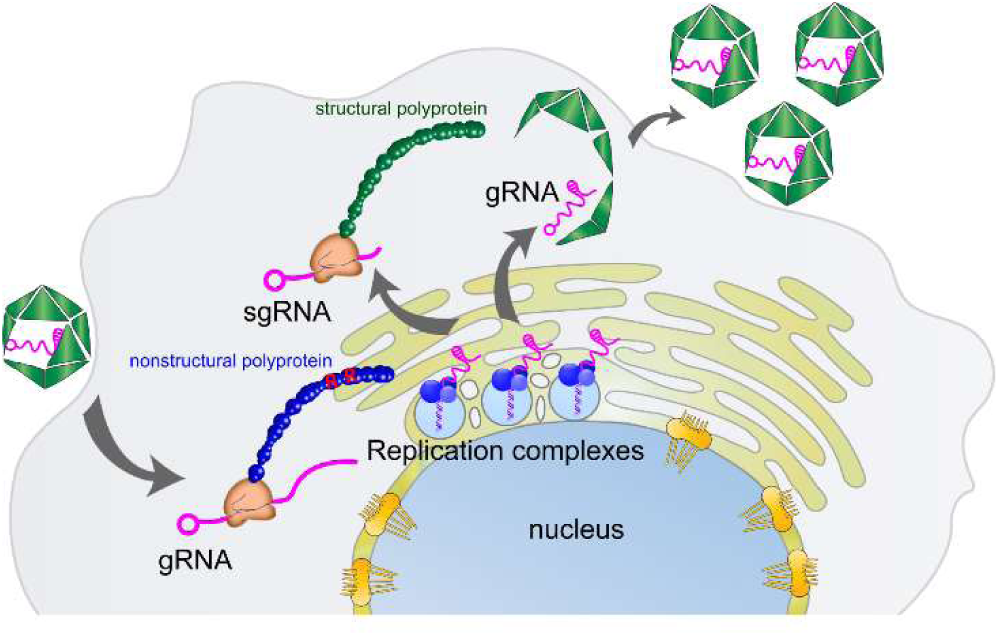
**The mechanism of NTD-directed formation of astrovirus replication complex.**

Perinuclear localization of replication complexes has been reported for several RNA viruses. In particular, perinuclear ER-specific replication is a hallmark of the replication of brome mosaic virus ^33,41^, flaviviruses ^42,43^, and several other virus families. The mechanisms of perinuclear ER targeting represent a diverse set of approaches based on molecular mimicry of ER-residing proteins and recruitment of related ER-associated proteins ^44^. We demonstrate a similar strategy employed by astroviruses thus expanding the list of viral ER hijackers. An outstanding question is how host factors cooperate with virus-encoded di-arginine motifs to construct the ER-derived replication organelle and how virus assembly and maturation are linked to the modified cellular membranes.

Altogether, our findings provide insight into astrovirus replication and the involvement of the N-terminal nonstructural protein in the correct positioning of RCs. Future studies will address the exact host components involved in the formation of active replication organelles and describe the topology and functionality of viral nonstructural proteins. Understanding virus replication mechanisms will potentially lead to the development of future therapeutics.

## Materials and Methods

### Cells

BSR cells (single clone of BHK-21 cells) were maintained at 37 °C in DMEM supplemented with 10% fetal bovine serum (FBS), 1 mM L-glutamine and antibiotics. Caco2 and Huh7.5.1 cells (Apath, Brooklyn, NY) were maintained in the same media supplemented with non-essential amino acids. All cells were tested mycoplasma negative throughout the work (MycoAlert Mycoplasma Detection Kit, Lonza).

### Virus strains

HAstV1 (pAVIC1, L23513.1), MLB1 (pMLB1, ON398705) and MLB2 (pMLB2, ON398706) were derived from reverse genetics clones. VA1 astrovirus was kindly provided by David Wang (University of St Louis, USA) and clinical HAstV4 was kindly provided by Susana Guix (University of Barcelona, Spain).

### Plasmids and cloning

Reverse genetics and replicons of the human astroviruses MLB2 ^24^ and HAstV1 ^7,45^ were previously described. HA-tagged human astroviruses (MLB1 and HAstV1) were generated by site-directed mutagenesis. The coding sequence of the HA-tag was inserted in the nsP1a as shown in Fig 2A. The resulting infectious clones were designated as HAstV1-HA and MLB2-HA.

For mammalian expression of the MLB2 and HAstV1 nsP1a N-terminal domain, the coding sequence of the HA-tagged N-terminal domain from corresponding HA-tagged virus was inserted into vector pGAC-PM ^7^ using *Afl*II and *Pac*I restriction sites. The resulting constructs – designated pCAG-HAstV1-NTD-HA and pCAG-MLB2-NTD-HA – were confirmed by sequencing. All mutations were introduced using site-directed mutagenesis and confirmed by sequencing.

EV7 replicon was generated by insertion of mCherry coding sequence flanked by 3Cpro and 2A cleavage sites between nucleotides 794 and 3323 of pT7-EV7 plasmid (AF465516) ^46^.

### Recovery of recombinant viruses from T7 RNAs

The linearised infectious clones of HAstV1 (pAVIC1, L23513.1) and MLB2 (pMLB2, ON398706) were used to produce capped T7 RNA transcripts using T7 mMESSAGE mMACHINE Transcription kit (ThermoFischer, AM1344) according to the manufacturer’s instructions. Recombinant viruses were recovered from T7 transcribed RNA using electroporation of Huh7.5.1 cells (MLB2) or BSR cells (HAstV1) in PBS at 800 V and 25 µF using a Bio-Rad Gene Pulser Xcell electroporation system. For HAstV1 passaging, the collected supernatant was treated with 10 µg mL^−1^ trypsin (Type IX, Sigma, #T0303) for 30 min at 37 °C, diluted 5 times with serum-free media, and used for infection of Caco2 cells. The passaging of MLB2 was performed on Huh7.5.1 cells in the absence of trypsin. Recombinant (HAstV1, MLB1, MLB2) and clinically isolated (HAstV4, VA1) viral stocks were titrated using immunofluorescence-based detection with 8E7 (HAstV1, HAstV4) or custom polyclonal antibodies against capsid proteins (MLB1, MLB2, VA1) ^7,24^.

### Virus growth curves

Multistep growth curves were performed using an MOI of 0.1, with infections performed in triplicates. Media-derived samples were collected in equal amounts at 0, 24, 48, 72 and 96 hpi and titrated. The titers were determined as infectious units per ml (IU/ml).

### Electroporation of plasmid DNA

To analyze overexpressed proteins, electroporation of Huh7.5.1 cells was performed using 3 µg of plasmid DNA in full media at 220 V and 975 µF using a Bio-Rad Gene Pulser, in the presence of 5 mM NaBes (N,N-Bis(2-hydroxyethyl)-2-aminoethanesulfonic acid sodium salt) and 0.2 mg/ml salmon sperm carrier DNA. After electroporation, cells were split between immunofluorescent and western blotting analyses and incubated for 24 hours.

### SDS-PAGE and immunoblotting

Proteins were resolved on a 12 or 15% SDS-PAGE gel before being transferred to 0.2 µm nitrocellulose membrane and blocked in 4% Marvel milk powder in PBS. Immunoblotting with Astrovirus 8E7 antibody (Santa Cruz Biotechnology, sc-53559, 1:1000), MLB1 anti-capsid (custom ^24^, 1:500), anti-HA (Abcam, ab130275, 1:3000) and anti-tubulin antibody (Abcam, ab6160, 1:1000) was followed by Licor IRDye 800 and 680 secondary antibodies (1:3000). Immunoblots were imaged on a LI-COR ODYSSEY CLx imager and analyzed using Image Studio version 5.2.

### Immunofluorescence

Infected (Fig 2-5) or electroporated (Fig 6) cells were grown on IBIDI wells, then washed with PBS and fixed with 10% formaldehyde in PHEM buffer (60 mM Pipes, 25 mM Hepes, 10 mM EGTA, 2 mM MgCl_2_, pH 7.0) for 15 min. Fixed cells were then washed with PBS and permeabilized with 0.1% Triton X100 or 0.1% saponin for 10 min, followed by blocking in 2% goat serum in PBS for 1 hour. For lamin staining, cells were fixed in 20% cold methanol for 15 mins. Blocked cells were stained with the following primary antibodies: MLB1 anti-capsid (1:300), Astrovirus 8E7, anti-dsRNA IgG2a (Scicons J2, 10010500, 1:250), anti-calnexin (Merck, MAB3126), anti-HA (Abcam, ab130275), anti-lamin A+C (Abcam, ab133256, 1:1000), followed by staining with Alexa 488- or 594-conjugated secondary antibody (Thermo Fisher, 1:1000). Nuclei were counter-stained with Hoechst. All confocal images are single-plane images taken with a Leica SP5 Confocal Microscope using a water-immersion 63× objective.

### MLB2 and HAstV1 replicon assay

Linearized replicon-encoding plasmids were used to generate T7 RNAs using T7 mMESSAGE mMACHINE Transcription kit (ThermoFischer, AM1344) according to the manufacturer’s instructions, purified using Zymo RNA Clean & Concentrator kit and quantified. Huh7.5.1 cells were transfected in triplicate with Lipofectamine 2000 reagent (Invitrogen), using previously described reverse transfection protocol ^24^. Three independent experiments, each in triplicate, were performed to confirm the reproducibility of the results.

### Statistical analyses

Data were graphed and analyzed using GraphPad Prism and MS Excel. Where appropriate, data were analyzed using one-way, two-way ANOVA, Student’s t-test or two-tailed Mann-Whitney test. Significance values are shown as \*\*\*\**p* < 0.0001, \*\*\**p* < 0.001, \*\**p* < 0.01, \**p* < 0.05, ns – non-significant.

## Acknowledgments

This work was funded by a Sir Henry Dale Fellowship (220620/Z/20/Z) from the Wellcome Trust and the Royal Society, an Isaac Newton Trust/Wellcome Trust ISSF/University of Cambridge Joint Research Grant and MRC project grant (MR/T000376/1) to V.L. D.N. is supported by the Department of Pathology studentship from Elisabeth Mann Fund. We thank Jean Thompson for excellent technical assistance and the Cambridge NIHR BRC Cell Phenotyping Hub for access to confocal microscopy.

